# Stimulus expectations do not modulate visual event-related potentials in probabilistic cueing designs

**DOI:** 10.1101/2023.04.05.535778

**Authors:** Carla den Ouden, Andong Zhou, Vinay Mepani, Gyula Kovács, Rufin Vogels, Daniel Feuerriegel

**Affiliations:** Melbourne School of Psychological Sciences, The University of Melbourne; Institute of Psychology, Friedrich Schiller University Jena, Jena, Germany; Laboratorium voor Neuro-en Psychofysiologie, Department of Neurosciences, KU Leuven, Leuven, Belgium

**Keywords:** EEG, ERP, expectation, surprise, prediction

## Abstract

Humans and other animals can learn and exploit repeating patterns that occur within their environments. These learned patterns can be used to form expectations about future sensory events. Several influential predictive coding models have been proposed to explain how learned expectations influence the activity of stimulus-selective neurons in the visual system. These models specify reductions in neural response measures when expectations are fulfilled (termed expectation suppression) and increases following surprising sensory events. However, there is currently scant evidence for expectation suppression in the visual system when confounding factors are taken into account. Effects of surprise have been observed in blood oxygen level dependent (BOLD) signals, but not when using electrophysiological measures. To provide a strong test for expectation suppression and surprise effects we performed a predictive cueing experiment while recording electroencephalographic (EEG) data. Participants (n=48) learned cue-face associations during a training session and were then exposed to these cue-face pairs in a subsequent experiment. Using univariate analyses of face-evoked event-related potentials (ERPs) we did not observe any differences across expected (90% probability), neutral (50%) and surprising (10%) face conditions. Across these comparisons, Bayes factors consistently favoured the null hypothesis throughout the time-course of the stimulus-evoked response. When using multivariate pattern analysis we did not observe above-chance classification of expected and surprising face-evoked ERPs. By contrast, we found robust within– and across-trial stimulus repetition effects. Our findings do not support predictive coding-based accounts that specify reduced prediction error signalling when perceptual expectations are fulfilled. They instead highlight the utility of other types of predictive processing models that describe expectation-related phenomena in the visual system without recourse to prediction error signalling.

**Highlights:**

– We performed a probabilistic cueing experiment while recording EEG.
– We tested for effects of fulfilled expectations, surprise, and image repetition.
– No expectation-related effects were observed.
– Robust within– and across-trial repetition effects were found.
– We did not find support for predictive coding models of expectation effects.

## 1. Introduction

Humans and other animals can learn from recurring patterns of sensory input and form predictions about upcoming events. For example, after hearing a loud bark one might expect to see a dog and would be surprised upon seeing a cat instead. These so-called predictive processes can help us to detect unusual or novel events (Ulanovsky et al., 2003; Press et al., 2020), identify deviations from regular sequences of stimuli (Fiser & Aslin, 2002; Sherman et al., 2020) and make faster perceptual decisions when our expectations are fulfilled (Mulder et al., 2012; Gold & Stocker, 2017).

A plethora of theories and models have been proposed to explain how predictive processes influence the activity of stimulus-selective neurons in the visual system (e.g., Rao & Ballard, 1999; Friston, 2005, 2010; Summerfield & de Lange, 2014; Spratling, 2017; Keller & Mrsic-Flogel, 2018; Hogendoorn & Burkitt, 2019; Press et al., 2020). Among these, a set of widely-influential hierarchical predictive coding models (Friston, 2005, 2010; Bastos et al., 2012; Clark, 2013; Summerfield & de Lange, 2014) have been developed based on the premise that sensory systems are continuously attempting to infer the underlying causes of incoming sensory signals. This is proposed to occur via generation and continuous updating of internal models of an organism’s sensory environment. Discrepancies between the internal model and sensory input are thought to be indexed by activity of prediction error neurons that are prevalent throughout the cortical hierarchy of the visual system (e.g., Friston, 2010). According to these predictive coding models, an organism can use past experience to update its internal model in anticipation of expected sensory input.

When an observed stimulus matches the organism’s expectations the degree of prediction error (and activity of prediction error neurons) should be less than when the stimulus is not expected (Summerfield & de Lange, 2014). Here, we describe prediction errors as indexing the *absolute* degree of discrepancy between expectations and sensory input with respect to a given feature space (in contrast to signed reward prediction errors as defined in Schultz, 2016). We define expectations as predictions to see certain stimuli (or stimulus features) that are formed *prior* to the appearance of the stimulus (in contrast to e.g., Rao & Ballard, 1999, see Andersen & Chemero, 2013).

The diversity of predictive coding implementations (e.g., Bastos et al., 2012; Spratling, 2017; Keller & Mrsic-Flogel, 2018) makes it difficult to derive specific model predictions that are both i.) shared across models and ii.) unique to predictive coding accounts (see Cao, 2020; Milkowski & Litwin, 2022). However, there is one hypothesis that is relatively consistent across the models described above: that neural response measures should inversely scale with the subjective appearance probability of a presented stimulus (Walsh et al., 2020; Feuerriegel et al., 2021a). In the visual system this hypothesis has been tested by entraining expectations to see particular stimuli in certain contexts. For example, this is achieved by consistently pairing a visual stimulus with a preceding cue (in predictive cueing designs, e.g., Egner et al., 2010) or manipulating the frequency with which different stimuli are presented (e.g., Kimura et al., 2009; Bell et al., 2016). Stimuli that conform to learned expectations are termed expected stimuli, and those that violate such expectations are termed surprising stimuli. A subset of studies also presented conditions whereby two or more different stimuli are each expected to appear with equal probability (termed neutral stimuli, for further definition see Arnal & Giraud, 2012; Feuerriegel et al., 2021a). We note that the terms ‘expected’, ‘surprising’ and ‘neutral’ are defined here in ways that are consistent with how these terms have been used in closely related work (e.g., Egner et al., 2010; Rahnev et al., 2011; Amado et al., 2016; Feuerriegel et al., 2018a; Dunovan & Wheeler, 2018, for further discussion see Feuerriegel et al., 2021a). These terms may not cleanly map onto concepts with similar names in other predictive processing frameworks, for example those that describe quantitative measures of surprise (e.g., Baldi & Itti, 2010; Loued-Khenissi & Preuschoff, 2020). Researchers have reported smaller fMRI BOLD signals (Summerfield et al., 2008; Egner et al., 2010; Grotheer & Kovács, 2015) and firing rates (Meyer & Olson, 2011; Bell et al., 2016) for expected compared to surprising stimuli.

These effects have been collectively called expectation suppression (ES; Summerfield et al., 2008; Todorovic & de Lange, 2012). According to predictive coding accounts, ES is assumed to reflect suppressed, silenced or dampened responses when one’s expectations are fulfilled compared to when there are no consistent expectations to see a particular stimulus (i.e., an expected-neutral condition difference, Summerfield et al., 2008; Egner et al., 2010). For example, ES would lead to smaller neural response measures when a single stimulus is cued to appear with high probability (an expected stimulus) compared to when two different stimuli could each be expected to appear with 50% probability (a neutral stimulus). These effects are thought to be analogous to effects of expectation violation whereby surprising stimuli (e.g., low appearance probability) evoke larger prediction error responses than expected or neutral conditions (Egner et al., 2010).

In our recent review (Feuerriegel et al., 2021a) we reported that, for fMRI BOLD signals in the visual system, there is actually limited and inconsistent evidence for ES defined as an expected-neutral difference. BOLD signal differences between expected and surprising stimuli were apparently due to surprise-related increases rather than any suppression associated with expectation fulfilment (e.g., Summerfield & Koechlin, 2008; Amado et al., 2016).

In addition, multiple confounding factors were identified in Feuerriegel et al. (2021a) that may mimic expectation effects in commonly used experiment designs. One prominent confound is effects of repetition suppression and adaptation, whereby stimuli that are identical (or similar) to those that were recently encountered evoke reduced neural response magnitudes (Grill-Spector et al., 2006; Solomon & Kohn, 2014; Vogels, 2016). These effects are dissociable from those of cued probabilistic expectations (Kaliukhovich & Vogels, 2011, 2014; Larsson & Smith, 2012; Grotheer & Kovács, 2015; Vinken et al., 2018; Tang et al., 2018; Feuerriegel et al., 2018a; Solomon et al., 2021). Importantly, adaptation effects can extend across experimental trials and intervening stimulus presentations (Bayliss & Rolls, 1987; Kaliukhovich & Vogels, 2014; Henson, 2016; Fritsche et al., 2022). Differences in the novelty of a stimulus (operationalised as the number of times a stimulus has been encountered in an experiment) can also lead to larger responses for relatively novel stimuli (Xiang & Brown, 1998; Mur et al., 2010; Manahova et al., 2018). Both adaptation and novelty effects are particularly problematic in experiments that entrain expectations by manipulating the presentation frequency of different stimuli (e.g., Bastos et al., 2020). Here, we note that some confounding factors have been described as consequences of biased expectations, such as repetition suppression (Summerfield et al., 2008; Auksztulewicz & Friston, 2016). If these descriptions are accurate, then expectation-related effects should still be observed even when controlling for these confounds.

When these confounding factors are taken into account, it appears that surprise-related effects on BOLD signals in predictive cueing designs (e.g., Amado et al., 2016) have not replicated when using electrophysiological measures, such as firing rates and local field potential amplitudes in macaques (e.g., Kaliukhovich & Vogels, 2011; Vinken et al., 2018; Solomon et al., 2021) and event-related potentials/fields (ERPs/ERFs) in humans (Kok et al., 2017; Rungratsameetaweemana et al., 2018; Solomon et al., 2021; reviewed in Feuerriegel et al., 2021a). Studies that did report expected-surprising differences found effects at frontal channels or topographies consistent with effects on the parietal P3 ERP component rather than sources in visual cortex (Summerfield et al., 2011; Hall et al., 2018; Feuerriegel et al., 2018a). Solomon et al. (2021) only reported differential ERPs to surprising stimuli when they were designated as targets in a go/no-go task. These effects resembled ERP components associated with target detection in perceptual decision tasks (e.g., Loughnane et al., 2016). ES and surprise effects have been observed in macaques during predictive cueing and statistical learning experiments that involved weeks of sequence learning prior to recording (Meyer & Olson, 2011; Meyer et al., 2014; Ramachandran et al., 2017; Schwiedrzik & Freiwald, 2017; Kaposvari et al., 2018; Esmailpour et al., 2022), however these effects have not been consistently replicated in human studies whereby sequences were learned over 1-2 testing sessions (e.g., Manahova et al., 2018; Zhou et al., 2020; reviewed in Feuerriegel et al., 2021a).

It is possible that expectation effects are more subtle than previously assumed and may require larger samples than what is typical of existing work in humans (e.g., n=23 in Kok et al., 2017; discussed in Feuerriegel et al., 2021a). In addition, participants could not form expectations for specific images in Rungratsameetaweemana et al. (2018) and expectation-related effects may be most prominent (or detectable) when there are specific expectations for a combination of low– and high-level features.

Providing strong tests for expectation effects on electrophysiological measures is important because prediction error signalling mechanisms are proposed to be directly related to changes in ERPs, firing rates and local field potentials (e.g., Friston, 2005; Bastos et al., 2012). By contrast, effects on fMRI BOLD signals are more difficult to interpret because they can arise from additional consequences of surprise, such as increased pupil dilation (O’Reilly et al., 2013; Richter & de Lange, 2019, discussed in Feuerriegel et al., 2021a).

Here, we tested for ES and surprise effects in humans using a larger (n=48) sample than previous electrophysiological studies while accounting for relevant confounding factors identified in Feuerriegel et al. (2021a). We used a predictive cueing design whereby a cue signalled the appearance probability of a subsequent face image, ranging between 10% (surprising), 50% (neutral) and 90% (expected). We compared ERPs evoked by expected, neutral and surprising faces using mass-univariate analyses to characterise expectation effects across the entire time-course of the stimulus-evoked response. We also tested for expected-surprising differences that are broadly distributed across the scalp using multivariate pattern analysis (MVPA). In each cueing condition only two different face images could potentially appear, meaning that participants could form expectations for specific low– and high-level stimulus features. In our design, expected and surprising faces were task-relevant but did not require an immediate decision or motor response. This allowed us to avoid ERP effects related to target detection or perceptual decision processes (e.g., O’Connell et al., 2012) that produced apparent surprise responses in one experiment of Solomon et al. (2021).

We additionally included another neutral expectation condition in which one of four face images could each be expected to appear with 25% probability (here termed the 25% neutral condition). This allowed us to test for a different expectation-related effect associated with the number of different stimuli that could potentially be expected to appear (predictability effects) identified in previous work (Pajani et al., 2017; Feuerriegel et al., 2018a; Rostalski et al., 2020). We also tested for stimulus repetition effects to compare their time-courses to effects of ES or surprise. If the time courses are different (as in Feuerriegel et al., 2018a) then this would further support the view that repetition suppression and ES are distinct phenomena (contrary to e.g., Summerfield et al., 2008; Auksztulewicz & Friston, 2016).

## 2. Method

### 2.1. Participants

Fifty participants were recruited for this experiment. Participants were fluent in English and had normal or corrected-to-normal vision. One participant was excluded because of inconsistent cue-face relationships used across the training session and experiment due to experimenter error. One additional participant was excluded because of a response keypad malfunction during the first two blocks of the training session. This left 48 participants for both behavioural and EEG data analyses (32 women, 16 men, 44 right-handed) aged between 18 and 35 (*M* = 22.0, *SD* = 2.9). This was a substantially larger sample than existing studies testing for electrophysiological expectation effects using predictive cueing designs (e.g., n=23 in Kok et al., 2017; n=17 in Rungratasemeetaweemana et al., 2018; n=22 in Solomon et al., 2021). As these studies did not report statistically significant expectation effects, we did not have an *a priori* effect size target for statistical power analyses. Instead, we aimed to maximise measurement precision for frequentist and Bayesian statistical tests. Here we note that, if the activity of superficial pyramidal neurons in visual cortex reflects prediction errors (Friston, 2005; Bastos et al., 2012), one might expect to observe moderate to large effects of stimulus expectations on EEG signals. Participants were reimbursed 45 AUD for their time. This study was approved by the Human Research Ethics Committee of the University of Melbourne.

### 2.2. Stimuli

Stimuli were presented at a distance of 80cm on a 27” Benq RL2755 LCD monitor (60Hz refresh rate) in a darkened room. Stimuli were presented using MATLAB and functions from PsychToolbox (Brainard, 1997; Kleiner et al., 2007). Code used for stimulus presentation will be available at osf.io/ahuc4/ at the time of publication. Cue stimuli included seven animal images from the Bank of Standardised Stimuli (Brodeur et al., 2014). Four female faces of neutral expression (denoted here as faces A/B/C/D) were also presented (as used in Rostalski et al., 2020). Faces were greyscale and centred with the fixation cross between the eyes. Face images were cropped so that only the face was shown on a black background, omitting any hair, clothing or jewellery. Example stimuli are displayed in Figure 1. Face stimuli were used because previous experiments presenting faces have reported surprise-related effects using fMRI (e.g., Summerfield et al., 2008; Amado et al., 2016) and in visual oddball designs using EEG (e.g., Feuerriegel et al., 2021b). The first face that appeared in each trial subtended 5.87 * 5.30 degrees of visual angle. The second face image in each trial was 20% smaller to increase the difficulty of the face matching task (described below).

**Figure 1.**
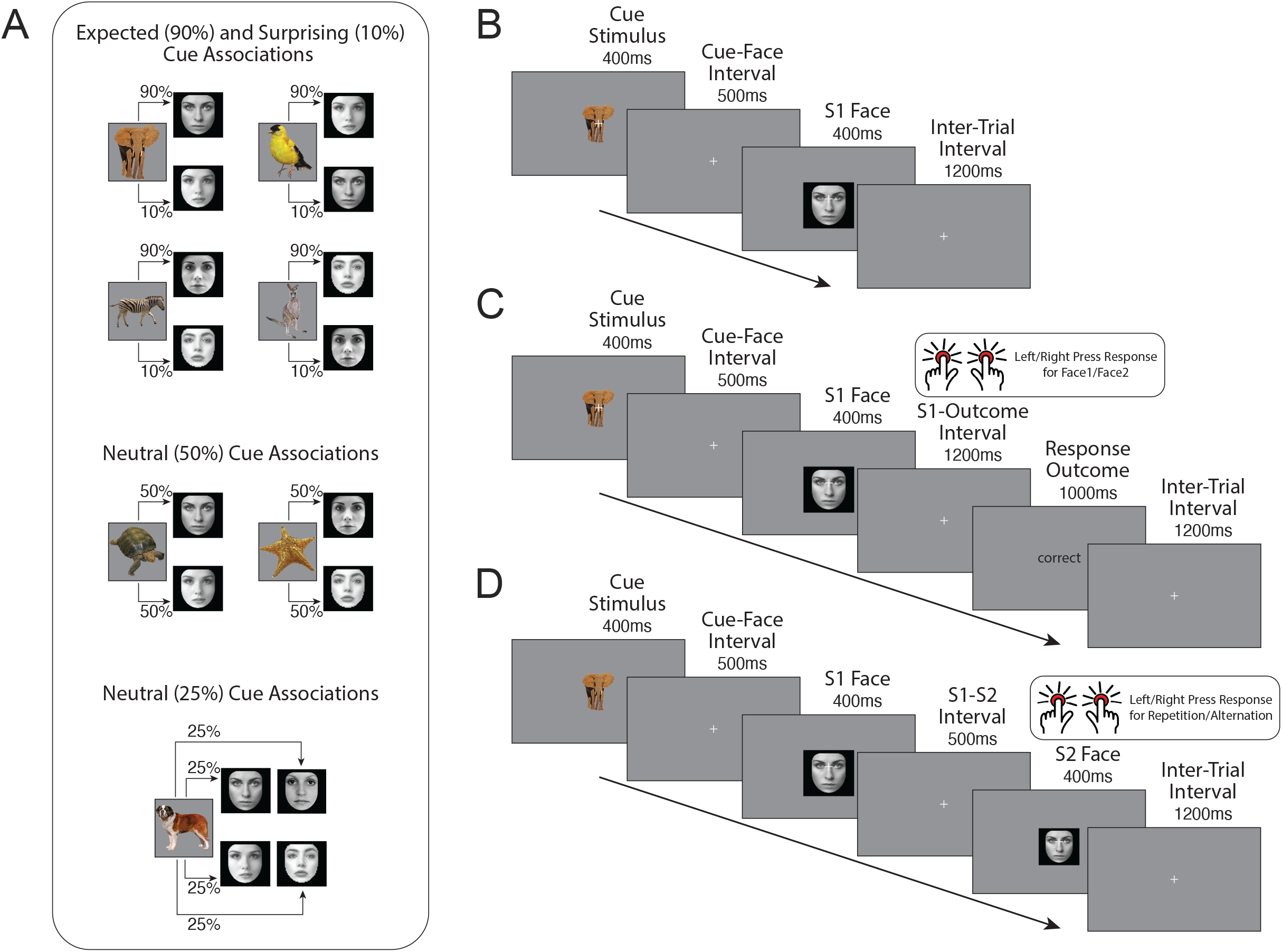
Cue-face associations and trial diagrams. A) An example of cue-face associations for a participant. Each animal image cue predicted the subsequent appearance of one of two face images with 90% probability (expected faces), 10% probability (surprising faces) or 50% probability (50% neutral faces). An additional cue predicted one of four faces to each appear with 25% probability (25% neutral faces). B) An example trial in the training session whereby participants were passively exposed to cue-face pairs. C) An example trial from the training session face identification task. Participants were required to press one of two keys depending on the face identity presented in each trial. For the 25% neutral cue condition, participants were instructed to press keys in response to two of the face identities, and not to respond if one of the other two faces were presented. D) An example trial from the experiment. Participants were required to press one of two keys based on whether the second (S2) face presented in the trial was the same or a different identity to the first (S1) face. Neither the animal cues or the S1 face identity were predictive of the S2 face being a repetition or alternation. All face images shown here are subject to a Pixabay license (https://pixabay.com/hu/service/license/). These images were not part of the actual stimulus set and we do not have permission to publish the original images. All images in this figure have been processed in the same way as the original stimuli.

### 2.3. Procedure

Prior to the experiment the EEG cap was set up and participants completed a training session to learn the probabilistic cue-face associations. Seven different cues were presented, which each cued the appearance probabilities of subsequently presented face images A/B/C/D. The first cue indicated that Face A would appear with 90% probability (an expected stimulus), and Face B with 10% probability (a surprising stimulus). The second cue instead indicated Face B would appear with 90% probability and Face A with 10% probability. For the third and fourth cues, the same probabilities were indicated in relation to Faces C and D. The fifth cue indicated that either Faces A or B could each appear with 50% probability (50% neutral stimuli), and the sixth cue indicated that faces C or D could each appear with 50% probability. The seventh cue indicated that Faces A, B, C, or D could each appear with 25% probability (25% neutral stimuli). These neutral conditions were included as comparison conditions to isolate and quantify ES and surprise effects. The 25% neutral condition was also included to test for effects of face image predictability (Rostalski et al., 2020) and to provide an additional comparison condition used to test for ES (discussed in Feuerriegel et al., 2021a). Examples of cue-face associations for a participant are shown in Figure 1A. Animal images allocated to each cue type were counterbalanced across participants.

During training, each of the seven cue-face associations was introduced in separate sections. At the beginning of each section participants were shown the animal cue image. They were then shown the associated faces and explicitly told the appearance probability of each face following the cue. Following this, participants passively viewed a block of trials during which a cue was presented for 400ms, and after a 500ms interval a face image was presented for 400ms. Following the offset of this face a fixation cross was presented for 1,200ms before the start of the following trial (Figure 1B). Faces followed the cues according to the instructed appearance probabilities. For example, for the cue that indicates 90% probability for Face A and 10% for Face B, Face A was presented nine times within the block and Face B was presented once. These blocks included ten trials each for cues 1-6 and 20 trials for the seventh cue.

After the presentation of the first two cues, participants then completed a face identification task. In each trial of this task, one of the two cues was presented for 400ms followed by a face image for 400ms after a 500ms interval (Figure 1C). Participants were required to press keys 1 or 3 on a TESORO Tizona numpad (1000Hz polling rate) using their left or right index finger depending on the face identity shown (A or B). In half of trials the first cue was presented, and in the other half the second cue was presented (randomly interleaved). During the task the overall probabilities (averaged across cues) of Faces A and B appearing were 50%. This task was repeated following the introduction of the third and fourth cues (presenting faces C and D), the fifth cue (Faces A/B) and the sixth cue (Faces C/D). After instructions relating to the seventh cue, participants were instructed to respond to only two of the four faces (A and C) using left/right index fingers and to not respond when the other faces appeared. Following their response, participants were provided with feedback regarding the accuracy of their choice (‘correct’ or ‘error’) if their response was made within 1200ms following face onset. If a response was not made in this window the feedback ‘too slow’ was presented. If responses were made prior to face onset the feedback ‘too fast’ was presented. Each face identification task block included 60 trials. This task served to further strengthen the learned cue-face associations and allowed us to assess whether participants had learned these associations based on task performance across conditions. The training session lasted approximately twenty minutes.

Right after the training session participants began the experiment. We used a probabilistic cueing design that minimises confounds related to stimulus repetition and recency that are prominent in some other designs (Feuerriegel et al., 2021a). In each trial (depicted in Figure 1D) an animal cue was first presented for 400ms. After a 500ms interstimulus interval a face image (termed the S1 face) was presented for 400ms. Following another 500ms interstimulus interval, the second face in the trial (S2 face) was presented for 400ms. After S2 face offset there was a 1200ms inter-trial interval during which a fixation cross was displayed. Participants completed a face matching task in which they were required to judge whether the S1 and S2 face images were the same (repetitions) or different (alternations) by pressing keys 1 and 3 on the numpad with their left and right index fingers. Key assignments were counterbalanced across participants. Participants were required to make a response within 1200ms after S2 face onset. If a response was not made during this window, the feedback ‘too slow’ was presented for 1,000ms. If responses were made prior to S2 face onset then ‘too fast’ feedback was presented. This task ensured that the S1 faces predicted by each cue were task-relevant, but did not require an immediate perceptual decision and keypress response (see also Kok et al., 2017).

Prior to the experiment participants completed practice trials until they were confident in their ability to perform the task. Practice trials were identical to those in the experiment except that participants were provided with feedback (‘correct’, ‘error’, ‘too fast’, ‘too slow’) for 1,000ms after the response window following S2 face onset in each trial. A fixation cross was displayed for 800ms between the offset of task feedback and the presentation of the cue in the subsequent trial.

There were 70 trials per block, consisting of 36 expected faces, four surprising faces, twenty 50% neutral faces, and ten 25% neutral faces. Each face image was presented an equal number of times within each block. Participants completed 12 blocks including a total of 840 trials. At the end of each block participants were notified of their accuracy and mean RT for correct responses in that block. Participants had self-paced breaks between each block (minimum 10 seconds). The experiment (excluding breaks) lasted approximately 48 minutes.

### 2.4. Task Performance Analyses

To assess whether participants had learned the cue-face associations in the training session we derived measures of accuracy (proportion correct) and mean RTs for trials with correct responses. Analyses were performed in JASP v0.16.4 (JASP Core Team). We compared performance across expected (90%), 50% neutral and surprising (10%) faces. Accuracy was near ceiling for some conditions and was not normally distributed and so Wilcoxon signed-rank tests were used to test for differences in accuracy across conditions. Paired-samples t tests were used to test for differences in mean RTs. Bonferroni corrections were used to adjust the alpha level for multiple comparisons. Bayesian versions of each test were also run to derive Bayes factors in favour of the alternative hypothesis (Cauchy prior distribution, width 0.707, 1,000 samples drawn for signed-rank tests). Because the training task for 25% probability faces included 50% of trials where no response was required (resulting in a hybrid face identification and go/no-go task) we did not compare performance between this and other conditions.

Performance on the face matching task during the experiment was measured by calculating accuracy and mean RTs for trials with correct responses. We compared accuracy across conditions using Wilcoxon signed-rank tests with Bonferroni corrections as described above. We compared mean RTs across S1 face appearance probability conditions using a one-way repeated measures ANOVA. A Greenhouse-Geisser correction was used to account for sphericity violations. RTs were also compared across repeated and alternating face conditions using a paired-samples t test. Bayesian versions of each test were also run in JASP.

### 2.5. EEG Data Acquisition and Processing

We recorded EEG using a 64-channel Biosemi Active II system (Biosemi, The Netherlands) with a sampling rate of 512 Hz. Recordings were grounded using common mode sense and driven right leg electrodes (http://www.biosemi.com/faq/cms&drl.htm). We added six additional electrodes: two electrodes placed 1 cm from the outer canthi of each eye, and electrodes placed above and below the center of each eye.

We processed EEG data using EEGLab v2022.0 (Delorme & Makeig, 2004) in MATLAB (Mathworks). The EEG dataset and data processing and analysis code will be available at osf.io/ahuc4/ at the time of publication. First, we identified excessively noisy channels by visual inspection (mean number of bad channels = 0.5, range 0-3) and excluded these from average reference calculations and Independent Components Analysis (ICA). Sections with large amplitude artefacts were also manually identified and removed. We then low-pass filtered the data at 40 Hz (EEGLab Basic Finite Impulse Response Filter New, default settings), re-referenced the data to the average of all channels and removed one extra channel (AFz) to compensate for the rank deficiency caused by average referencing. We duplicated the dataset and additionally applied a 0.1 Hz high-pass filter (EEGLab Basic FIR Filter New, default settings) to improve stationarity for the ICA. The ICA was performed on the high-pass filtered dataset using the RunICA extended algorithm (Jung et al., 2000). We then copied the independent component information to the non high-pass filtered dataset (e.g., as done by Feuerriegel et al., 2018a). Independent components associated with blinks and saccades were identified and removed according to guidelines in Chaumon et al. (2015). After ICA, we interpolated previously removed noisy channels and AFz using the cleaned dataset (spherical spline interpolation). EEG data were then high-pass filtered at 0.1 Hz.

The resulting data were segmented from –100ms to 800ms relative to cue onset, S1 face onset and S2 face onset. Epochs were baseline-corrected using the prestimulus interval. Epochs containing amplitudes exceeding ±100µV from baseline at any of the 64 scalp channels, as well as epochs from trials with ‘too fast’ or ‘too slow’ responses to S2 faces, were rejected (cues: mean epochs retained = 786 out of 840, range 629-837, S1 faces: mean = 792, range 636-839, S2 faces: mean = 793, range 629-838). Statistics relating to numbers of epochs retained per condition are displayed in Supplementary Table S1.

### 2.6. Mass-Univariate Analyses of ERPs

As face-evoked ERP components are most prominent at bilateral parietal and parieto-occipital channels (e.g., Quek & Rossion, 2017; Feuerriegel et al., 2021b) we defined a parieto-occipital region of interest (ROI) covering electrodes P7, P8, P9, P10, PO7 and PO8. Trial-averaged ERPs in each condition were averaged across channels within this ROI prior to analyses. To provide a more comprehensive coverage of visual cortex we defined an additional occipital ROI covering electrodes Oz, O1, O2, POz, and lz. Data from this ROI was used in follow-up analyses (results displayed in the Supplementary Material).

To test for expectation effects across the time-course of the S1 face-evoked response we used mass-univariate analyses of ERPs. Paired-samples t tests were performed at each time point relative to stimulus onset. Cluster-based permutation tests based on the cluster mass statistic were used to account for multiple comparisons (Maris & Oostenveld, 2007; 461 comparisons per analysis, 10,000 permutation samples, cluster forming alpha = .01, family-wise alpha = .05) using functions from the Decision Decoding Toolbox v1.0.5 (Bode et al., 2019). For each permutation sample, condition labels were swapped for a random subset of participants and paired-samples t tests were performed at each time point. All t values corresponding to uncorrected p-values of < .01 were formed into clusters with any neighbouring such t values. Adjacent time points were considered temporal neighbours. The sum of the t values within each cluster is termed the ‘mass’ of that cluster. The largest cluster masses from each of the 10,000 permutation samples were used to estimate the null distribution. The masses of each cluster identified in the original dataset were then compared to the null distribution. The percentile ranking of each cluster mass value relative to the null distribution was used to derive the p-value for each cluster. This method provides control over the weak family-wise error rate while maintaining high sensitivity to detect effects (Maris and Oostenveld, 2007, Groppe et al., 2011).

We performed separate sets of mass-univariate analyses to compare ERPs across different pairs of conditions. This allowed us to separately test for each hypothesised effect. To test for combined effects of ES and surprise (as done in previous work, e.g., Feuerriegel et al., 2018a; Richter & de Lange, 2019), we compared ERPs following S1 faces in expected (90%) and surprising (10%) conditions. To test for ES effects we compared ERPs across expected and 50% neutral, as well as expected and 25% neutral conditions. To test for effects of surprise we compared ERPs across surprising and 50% neutral conditions. To test for effects of image predictability (Feuerriegel et al., 2018a; Rostalski et al., 2020) we compared ERPs across 50% and 25% neutral conditions, in which either two or four face images could be expected to appear, respectively. To derive estimates of evidence for the null and alternative hypotheses at each timepoint we calculated Bayes factors at each time point for each comparison using Bayesian paired-samples t tests as implemented in the BayesFactor toolbox v2.3.0 (Krekelberg, 2022, default settings).

To compare the time-courses of stimulus repetition and expectation effects (as done in Feuerriegel et al., 2018a) we additionally tested for effects of within– and across-trial image repetition using mass-univariate analyses as described above. To test for within-trial face repetition effects we compared ERPs evoked by S2 faces when preceded by the same face as compared to a different S1 face identity. To test for across-trial repetition effects we compared ERPs evoked by cue images based on whether the cue in the previous trial was the same or a different image. We also compared ERPs evoked by S1 faces depending on whether the S2 face in the previous trial was the same or a different face identity.

### 2.7. Multivariate Pattern Classification Analyses

The ROI-based mass-univariate analyses described above are sensitive to between-condition amplitude differences that are consistent across participants but are less sensitive to ERP pattern differences that vary across individuals or effects that occur outside of the predefined ROIs. To better test for such effects we used MVPA using support vector machine (SVM) classification as implemented in DDTBOX v1.0.5 (Bode et a., 2019) interfacing LIBSVM (Chang & Lin, 2011). To test for the combined influence of ES and surprise effects we trained classifiers to discriminate between expected and surprising S1 faces. These were preceded by the same cue images, meaning that late ERP differences across cue images could not contribute to above-chance classification performance (e.g., via effects on pre S1 face ERP baselines). Please note that here we did not classify face identity, but rather the expected or surprising status of the S1 faces. To compare the classification accuracy time-courses against within-trial repetition effects we also trained classifiers to discriminate between S2 faces based on whether they were preceded by the same face (repetition trials) or a different face image (alternation trials).

We used a sliding window approach whereby epochs were split into non-overlapping analysis time windows of 10ms in duration. Within each time window EEG amplitudes were averaged separately for each of the 64 scalp channels to create a spatial vector of brain activity (64 features) corresponding to each trial. In cases where one condition included more epochs than the other, a random subset of epochs was drawn from the former condition to balance epoch numbers. Data from each condition were then split into five subsets of epochs. SVM classifiers (cost parameter C = 1) were trained on the first four subsets (80% of the data) from each condition and subsequently tested using the remaining subset to derive a classification accuracy measure. This was repeated until each subset had been used once for testing. This 5-fold cross-validation procedure was then repeated another five times with different epochs randomly allocated to each subset each time to minimise drawing biases. The average classification accuracy across all cross-validation steps and analysis repetitions was taken as the estimate of classification performance for each participant. This procedure was repeated with randomly permuted condition labels assigned to each trial to derive an empirical chance distribution.

Classification performance using the original data was compared against permuted-labels classification performance at the group level using paired-samples t tests on data from each analysis time window. As we aimed to detect above-chance classification performance we used one-tailed tests. We used cluster-based permutation tests (settings as described above) to correct for multiple tests across analysis time windows.

## 3. Results

### 3.1. Training Session Task Performance

Accuracy and mean RTs for each cueing condition in the face identification task are plotted in Figure 2A-B. Participants achieved higher accuracy for expected compared to surprising faces, *Z* = 4.79, *p* < .001, BF_10_ > 2,600, and 50% neutral compared to surprising faces, *Z* = 5.28, *p* < .001, BF_10_ > 5,800 (Figure 2A). Participants were slightly more accurate for 50% neutral as compared to expected faces, *Z* = 2.65, *p* = .008, BF_10_ = 7.42. This is likely due to a block order effect. 50% neutral face training blocks were always presented after blocks of expected and surprising faces, meaning that participants were better practiced at the task when identifying neutral faces. RTs were faster for expected compared to both 50% neutral, *t*(47) = –4.22, *p* < .001, BF_10_ = 209.92, and surprising faces, *t*(47) = –6.25, *p* < .001, BF_10_ > 128,900 (Figure 2B).

**Figure 2.**
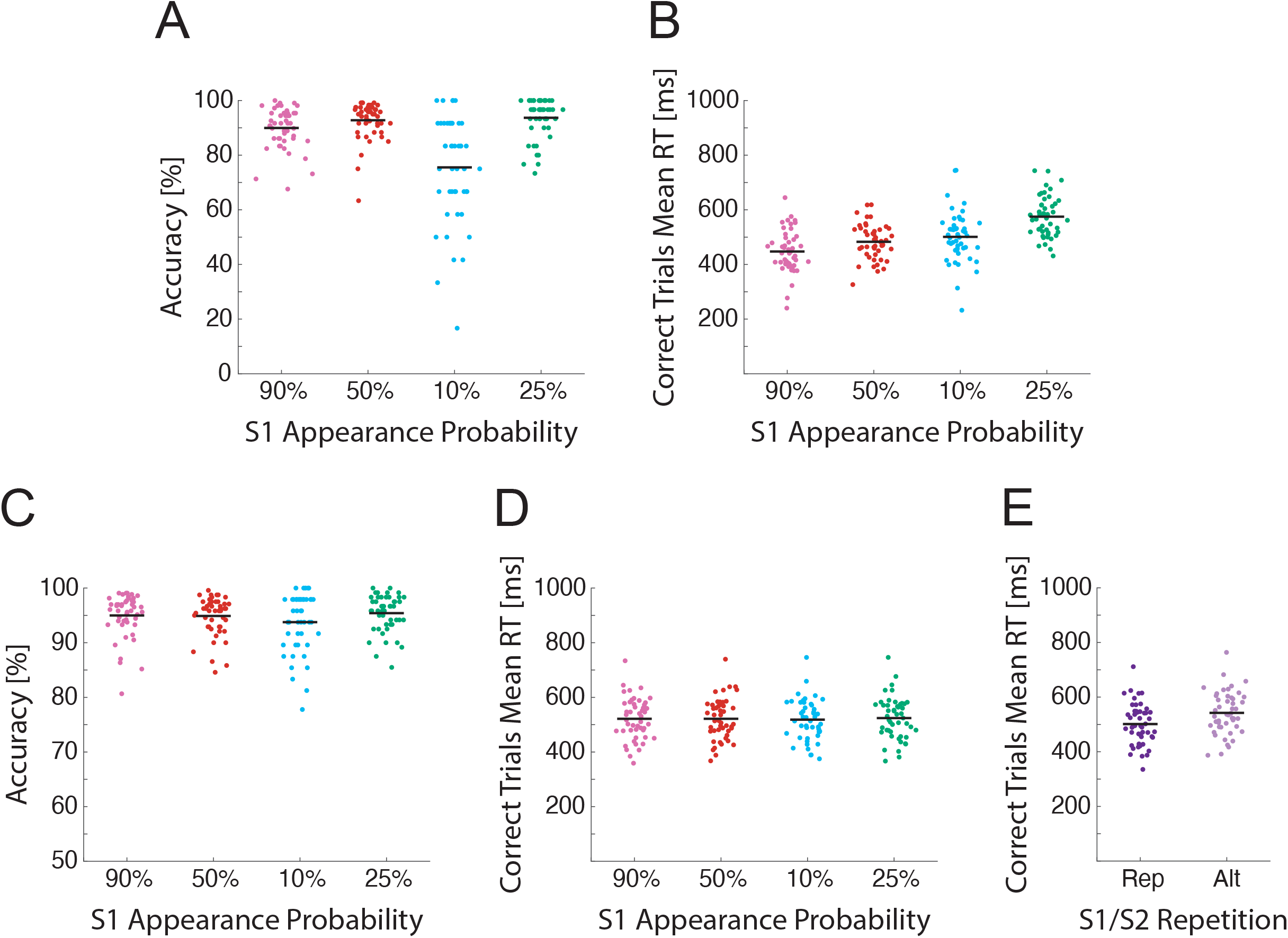
Task performance during the training session and experiment. A) Accuracy in the training session face identification task by cued stimulus appearance probability. Dots represent values for individual participants. Horizontal black lines denote group means. For the 25% probability condition accuracy refers to the proportion of target faces that were correctly identified. B) Mean RTs for correct trials. C) Accuracy in the face matching task by S1 face appearance probability condition. D) Mean RTs for correct trials by S1 face probability. E) Mean RTs for correct trials by S1-S2 repetition (Rep) or alternation (Alt) status.

However, we did not observe faster RTs for 50% neutral compared to surprising faces, *t*(47) = –1.56, *p* = .125, BF_10_ = 0.49. Higher accuracy and faster RTs for expected faces indicates that participants successfully learned the cue-face associations within the training session.

Please note that accuracy and mean RTs are plotted in Figure 2A-B also for 25% neutral faces requiring a keypress response. However due to the difference in task for this condition (with 50% of trials requiring no response) these were not compared with the other conditions.

### 3.2. Experiment Task Performance

During the experiment accuracy was generally high across all S1 face expectancy conditions for the S1-S2 face matching task (Figure 2C). Accuracy was slightly lower in trials with surprising as compared to expected S1 faces, *Z* = 2.06, *p* = .040, BF_10_ = 2.67, congruent with previous reports of impaired decision-making for stimuli immediately following surprising events (reviewed in Wessel, 2018). However this effect was not statistically significant when adjusting for multiple tests. Differences in accuracy were not observed when comparing other S1 face expectancy conditions, expected-50% neutral: *Z* = 0.55, *p* = .586, BF_10_ = 0.16, 50% neutral-surprising: *Z* = 1.89, *p* = .059, BF_10_ = 1.93, 50% neutral – 25% neutral: *Z* = –1.58, *p* = .115, BF_10_ = 0.67. Differences in mean RTs were not observed across expectancy conditions, *F*(2.15, 101.20) = 2.21, *p* = .112, BF_10_ = 0.37 (Figure 2D). RTs for repeated faces were faster than those for alternating faces, *t*(47) = –10.07, *p* < .001, BF_10_ = > 30^10^ (Figure 2E) consistent with Rostalski et al. (2020).

### 3.3. Expectation and Predictability Effects on S1 Face-Evoked ERPs

We did not observe statistically-significant ERP amplitude differences across any of the compared expectancy conditions. Group-averaged ERPs for each set of conditions, difference waves, standardised Cohen’s d effect size estimates (Cohen, 1988) and Bayes factors in favour of the alternative hypothesis are displayed in Figure 3. We did not observe differences between ERPs evoked by expected and surprising faces (Figure 3A), expected and 50% neutral faces (Figure 3B), expected and 25% neutral faces (Figure 3C), or 50% neutral and surprising faces (Figure 3D). Amplitude differences and standardised effect size point estimates were very small. Bayes factors generally provided evidence in favour of the null (values smaller than 1/3) across the time-course of the S1 face-evoked response.

**Figure 3.**
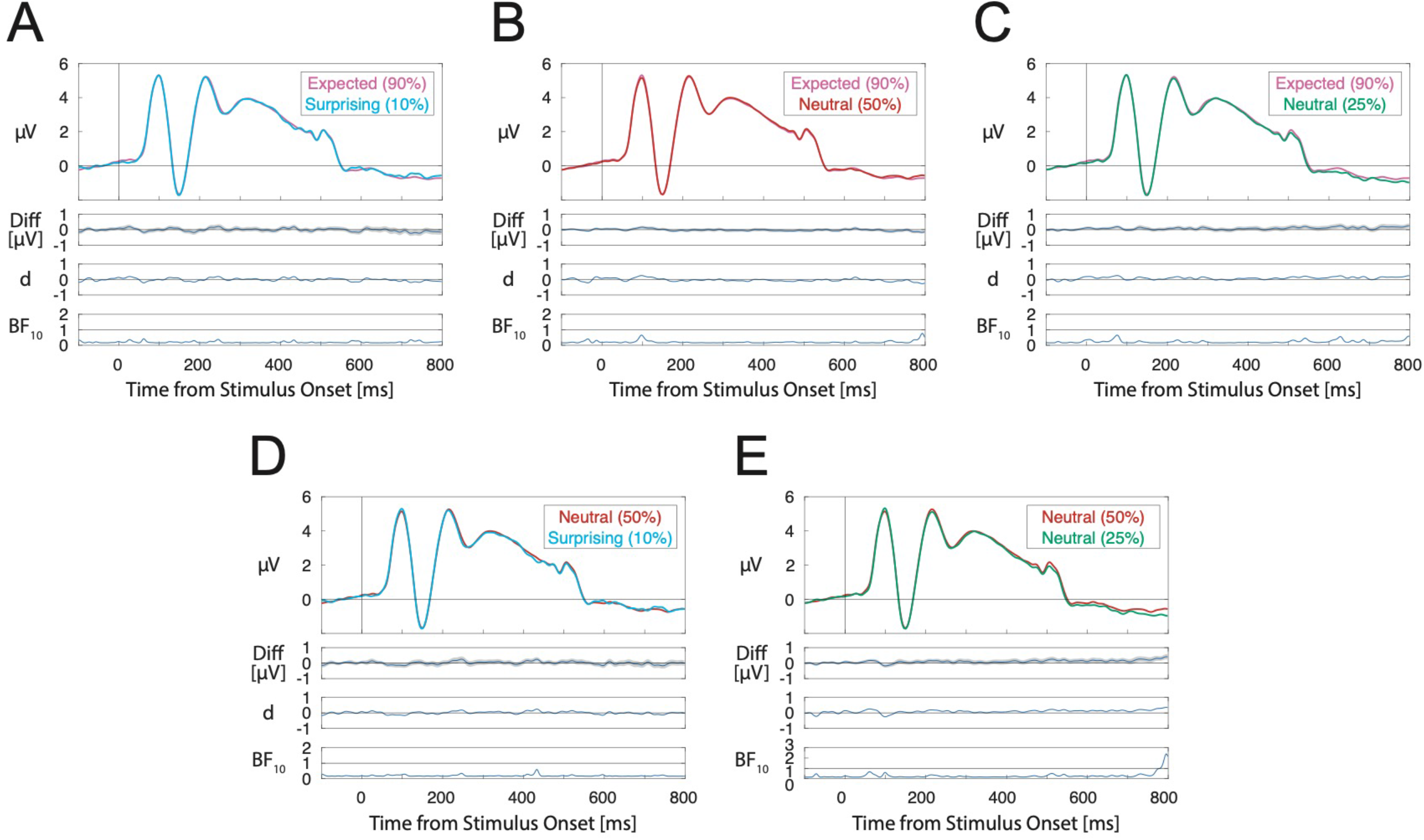
Group-averaged ERPs evoked by expected (90% appearance probability), surprising (10% probability), and neutral (50% and 25% probability) S1 faces. A) Expected – surprising ERP differences. B) Expected – 50% neutral differences. C) Expected – 25% neutral differences. D) 50% neutral – surprising differences. E) 50% neutral – 25% neutral differences. ERPs are averaged across channels within the parieto-occipital ROI including channels PO7/8, P7/8 and P9/10. ERPs for each pair of compared conditions are displayed along with difference waves (with shading denoting standard errors), Cohen’s d estimates and Bayes factors in favour of the alternative hypothesis.

We also compared ERPs across 50% and 25% neutral conditions to test for predictability effects (reported in Pajani et al., 2017; Feuerriegel et al., 2018a; Rostalski et al., 2020). Statistically-significant differences were not observed after correction for multiple comparisons. Bayes factors were in favour of the null hypothesis across most of the peristimulus time window (Figure 3E).

Results of analyses of ERPs at the occipital ROI (displayed in the Supplementary Material) were consistent with those described here.

### 3.4. Effects of Within-Trial Repetition on S2 Face-Evoked ERPs

Multiple within-trial face repetition effects were observed between 170-800ms from S2 face onset. Group-averaged ERPs for repeated and alternating S2 faces are displayed in Figure 4A. Scalp maps showing effect topographies are plotted in Figure 4B. This included an effect between 170-215ms (cluster mass = 90.58, critical cluster mass = 57.49, *p* = .044), an effect spanning 238-348ms (cluster mass = 409.79, *p* < .001) an effect spanning 379-453ms (cluster mass = 158.66, p = .009) and an effect spanning 754-800ms from S2 face onset (cluster mass = 92.79, p = .042). The timing and spatial distributions of these effects were broadly consistent with previously reported face repetition effects (e.g., Schweinberger & Neumann, 2016; Feuerriegel et al., 2018a, 2019).

**Figure 4.**
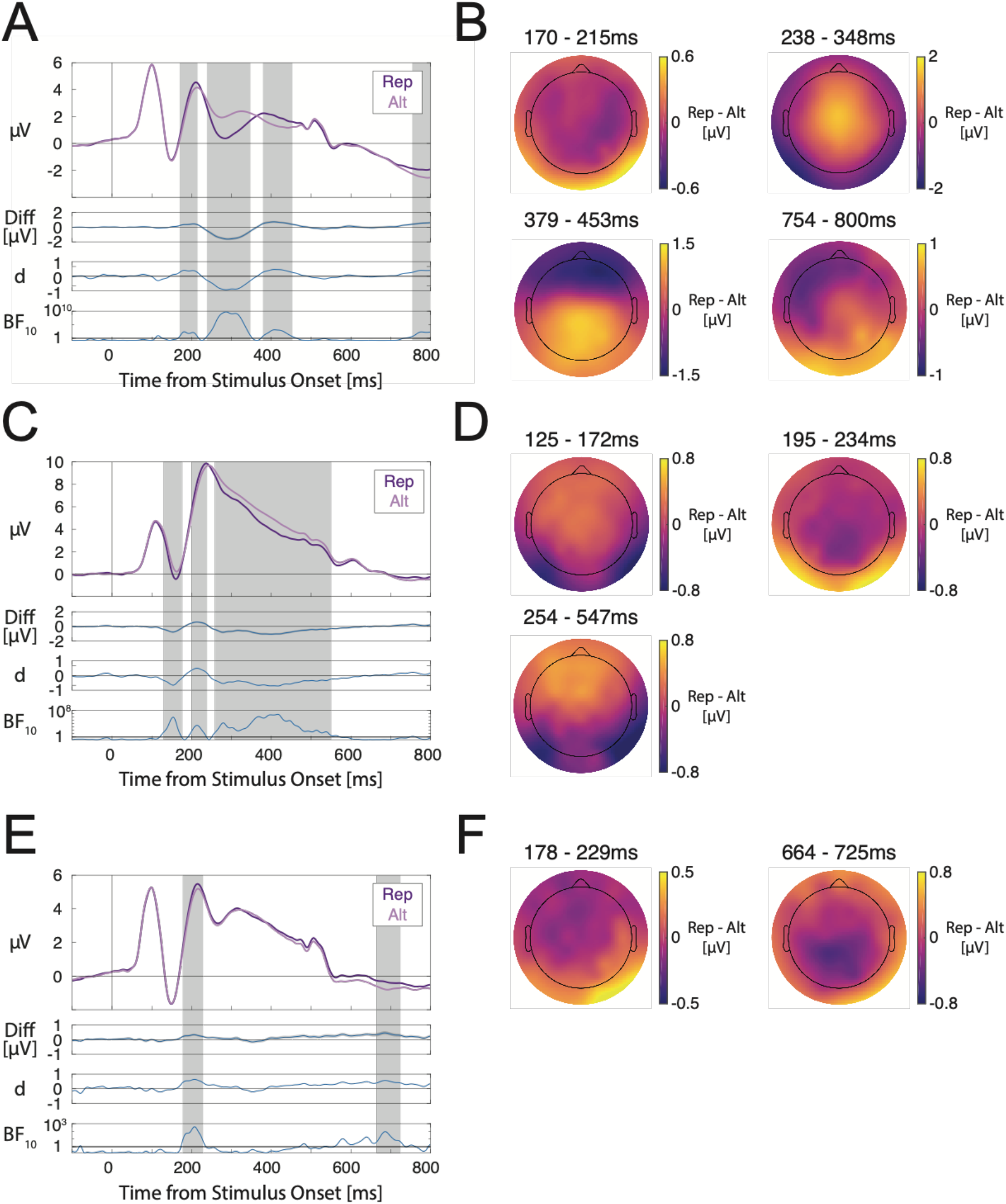
Within– and across-trial repetition effects for S2 faces, cues, and S1 faces. A) S2 face-evoked ERPs depending on whether the S2 face was the same as the S1 face identity (repetition/Rep) or a different face (alternation/Alt). B) Scalp maps of face repetition effects for S2 faces averaged over each statistically significant time window. C) Cue-evoked ERPs depending on whether the cue image was the same as the cue in the previous trial (Rep) or a different cue (Alt). D) Scalp maps of across-trial cue repetition effects averaged over each statistically significant time window. E) S1 face-evoked ERPs depending on whether the S2 face in the previous trial was the same (Rep) or a different identity (Alt). E) Scalp maps of across-trial S1 face repetition effects averaged over each statistically significant time window. ERPs are averaged across channels within the parieto-occipital ROI including channels PO7/8, P7/8 and P9/10. ERPs for each pair of compared conditions are displayed along with difference waves (with shading denoting standard errors), Cohen’s d estimates and Bayes factors in favour of the alternative hypothesis. Please note that Bayes factors are plotted on logarithmic scales due to the wide ranges of values across the time-course of the stimulus-evoked response. Grey shaded areas denote time windows of statistically significant differences.

### 3.5. Effects of Across-Trial Repetition on Cue– and S1 Face-Evoked ERPs

A number of across-trial cue image repetition effects were observed spanning 125-547ms from cue onset. Group–averaged ERPs evoked by cues that were either the same or different to the cue in the preceding trial are presented in Figure 4C. Cue repetition effects were observed between 125-172ms (cluster mass = 121.74, critical cluster mass = 47.50, *p* = .014), between 195-234ms (cluster mass = 87.22, *p* = .032) and between 254-547ms from cue stimulus onset (cluster mass = 769.71, *p* < .001). Topographies of effects (displayed in Figure 4D) resembled those observed for within-trial face repetition effects.

We also compared S1 face-evoked ERPs across conditions whereby the S2 face in the preceding trial was the same or a different face identity. Group-averaged ERPs are displayed in Figure 4E and scalp maps of effects are shown in Figure 4F. Across-trial repetition effects were observed between 178-229ms (cluster mass = 102.57, critical cluster mass = 39.88, *p* = .019) and between 664-725ms from S1 face onset (cluster mass = 105.78, *p* = .018).

### 3.6. Multivariate Pattern Classification Results

We used MVPA to test for patterns of single-trial ERPs that discriminate between expected (90% probability) and surprising (10%) S1 faces. S1 face expectancy could not be classified at above-chance levels at any time during the epoch (Figure 5A). We did not attempt to classify between other expectancy conditions as the preceding cue images systematically differed across these conditions within each participant dataset. In these cases, late cue image-specific ERPs can influence S1 face pre-stimulus baselines to produce spurious above-chance classification performance.

**Figure 5.**
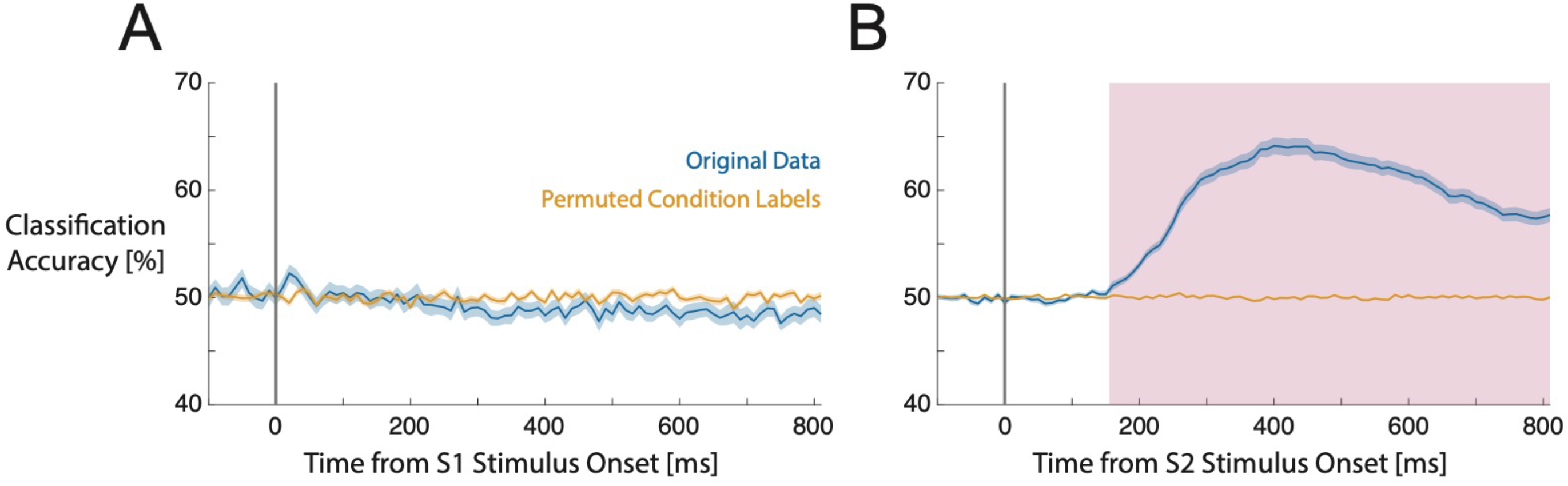
Multivariate pattern classification results. A) Classification accuracy for discriminating between expected (90% appearance probability) and surprising (10% probability) S1 faces. Blue lines display classification performance for the original data and orange lines show performance for the data with permuted condition labels (empirical chance distribution). Shaded regions denote standard errors. B) Classification accuracy for discriminating between repeated and alternating S2 faces. The magenta shaded area denotes the time window of statistically significant above-chance classification performance.

The within-trial repetition or alternation status of S2 faces could be classified at above-chance levels from 160 ms after S2 face onset until the end of the epoch (cluster mass = 948.72, critical cluster mass = 4.73, cluster p < .001).

## 4. Discussion

A set of influential predictive coding models specify that the response magnitudes of visual stimulus-selective neurons should inversely scale with the subjective appearance probability of a stimulus (Friston, 2005, 2010; Summerfield & de Lange, 2014; Walsh et al., 2020). To test this hypothesis, we entrained participants’ expectations to see particular face identities after certain cue images. Task performance results indicated that participants had learned these cue-face associations during a training session. We then presented the same cue-face associations in our experiment. Importantly, the cued faces were task-relevant but did not require an immediate decision or motor action. Despite analysing a large (n=48) sample we did not observe effects of either ES or surprise on face-evoked ERPs. Bayes factors instead signalled substantial support for the null hypothesis throughout the time-course of the face-evoked response (BF_10_ values of ∼0.2 for the majority of time points for each comparison). We also did not replicate image predictability effects observed in Feuerriegel et al. (2018a). However, we observed within– and across-trial image repetition effects for cue and face stimuli.

Our results, taken together with multiple null findings reported in earlier studies, indicate that expectations in predictive cueing designs are not sufficient to produce differences in stimulus-evoked electrophysiological responses within the human visual system. This is incongruent with models that specify reduced prediction error signalling when an observer’s expectations are fulfilled (e.g., Friston, 2005, 2010; Clark, 2013; Summerfield & de Lange, 2014) but is consistent with a growing body of electrophysiological work using predictive cueing designs that did not report effects of ES or surprise on responses of visual stimulus-selective neurons (Kaliukhovich & Vogels, 2011, 2014; Kok et al., 2017; Vinken et al., 2018; Rungratsameetaweemana et al., 2018; Solomon et al., 2021).

We additionally report that both within– and across-trial repetition effects can be readily observed in the absence of expectation effects. This is in line with most contemporary models of adaptation (e.g., Solomon & Kohn, 2014; Vogels, 2016) but is incongruent with predictive coding-based models that describe repetition suppression as a product of biased expectations toward previously-seen stimuli (e.g., Summerfield et al., 2008; Auksztulewicz & Friston, 2016).

### 4.1. Consequences of Cued Expectations on ERPs

We did not observe evidence for either ES or surprise effects on face-evoked ERPs. Bayes factors instead indicated evidence in favour of the null hypothesis throughout the peristimulus time window. Even when using MVPA, which can exploit distributed patterns of EEG signals across the scalp and ERP differences that are idiosyncratic to participants, we did not see above-chance classification accuracy when comparing expected and surprising faces.

Our findings are consistent with previous work using predictive cueing designs and electrophysiological recordings that did not report expectation effects (Kok et al., 2017; Rungratsameetaweemana et al., 2018; Kaliukhovich & Vogels, 2011; Vinken et al., 2018; Solomon et al., 2021; reviewed in Feuerriegel et al., 2021a). Although ERP differences between expected and surprising stimuli have been reported (Summerfield et al., 2011; Hall et al., 2018; Feuerriegel et al., 2018a) the topographies of these effects more closely indicate frontal sources or effects on the centro-parietal positivity ERP component (O’Connell et al., 2012; Twomey et al., 2015) rather than sources in the visual system. Tang et al. (2018) reported smaller visual N1 component peak amplitudes for surprising gratings, however this was likely a result of peak amplitude measurement biases due to a smaller number of trials per participant in the surprising condition (Thomas et al., 2004). Additionally, Solomon et al. (2021) reported expected-surprising differences only when surprising stimuli were targets, but the timing and topography of these effects closely resembled target detection-related ERP components observed in similar perceptual decision-making tasks (Loughnane et al., 2016). By contrast, in our design a perceptual decision was not required immediately following the expected and surprising S1 faces. Other effects of stimulus appearance probability outside of the visual system, such as those on P3 component amplitudes in oddball designs (Duncan-Johnson & Donchin, 1977; Polich & Margala, 1997), also appear to be absent when the critical stimuli do not prompt an immediate decision (Kimura et al., 2009; Feuerriegel et al., 2018b, 2021b; Amado & Kovács, 2016; File & Czigler, 2019).

At least three distinct possibilities arise from these findings. The first is that, in our study and in previous work, participants did not learn the predictive associations between cues and subsequently presented stimuli. We consider this unlikely because participants were explicitly told the cue-face contingencies in our experiment and demonstrated faster RTs for expected stimuli in the training session task (see also Kok et al., 2017; Rungratsameetaweeana et al., 2018; Vinken et al., 2018). During our experiment participants were further exposed to the expected cue-face pairs at least 100 times per cue. This would be expected to provide at least an equivalent opportunity to learn cue-stimulus associations as compared to existing fMRI predictive cueing designs (e.g., Egner et al., 2010).

The second possibility is that effects of fulfilled expectations and surprise are not expressed in neural activity that contributes to EEG and other electrophysiological measures, but are instead captured by changes in BOLD signals. Effects of surprise (but not ES) in predictive cueing designs have been identified and replicated across fMRI studies (e.g., Summerfield & Koechlin, 2008; Amado et al., 2016; reviewed in Feuerriegel et al., 2021a). However, this would run counter to the idea that prediction errors are signalled by cortical pyramidal neurons that typically contribute to invasive recordings and EEG/MEG (e.g., Friston, 2005; Bastos et al., 2012).

Notably, the inferences drawn from fMRI studies are complicated by additional effects of surprise (separate from prediction error signalling) that contribute to the BOLD response. For example, Richter & de Lange (2019) reported expected-surprising BOLD signal differences for object images only when participants completed an object categorisation task, and not during a task in which the objects were task-irrelevant (see also Larsson & Smith, 2012). In addition, they found that RT differences between expected and surprising conditions in the categorisation task correlated with BOLD signal differences across the same conditions in V1 and temporal occipital fusiform cortex. RTs are typically faster for expected stimuli (e.g., den Ouden et al., 2010; Mulder et al., 2012; Richter et al., 2018) and similar associations between RTs and BOLD signals in visual areas have been documented across decision-making tasks (Yarkoni et al., 2009; Mumford et al., 2023). Such associations are thought to reflect variation in the duration of decision-making processes that involve attention and visual working memory, termed a time-on-task effect (Yarkoni et al., 2009). Importantly, the probabilistic cueing studies that reported surprise effects on BOLD signals did not include RT as a regressor in their models. Such effects are also relevant to designs whereby participants detect a rare target that is distinct from the critical stimuli (e.g., inverted faces, Summerfield et al., 2008; Larsson & Smith, 2012; Feuerriegel et al., 2018a). This is because each stimulus requires a decision to determine that it is not a target, even if no motor response is required in target-absent trials (akin to a go/no-go task, Murphy et al., 2015). Time-on-task effects could be interpreted within predictive coding frameworks as reflecting the duration of prediction error signalling, where model convergence is slower for surprising stimuli. However, this would be incongruent with the finding that surprise effects were task-dependent in Richter and de Lange (2019).

Richter and de Lange (2019) also identified increased pupil dilation for surprising stimuli from ∼600 ms post stimulus onset, which correlated with the degree of expected-surprising BOLD signal differences in V1. Notably, this pupil dilation effect was not present when object stimuli were task-irrelevant. This can be understood as a secondary consequence of surprise, whereby increased pupil dilation leads to changes in retinal input and corresponding effects on low temporal resolution BOLD signals in visual cortex. In addition, BOLD signal increases for surprising stimuli have also been attributed to attentional capture once a stimulus is identified as unexpected (see Press et al., 2020; Alink & Blank, 2021). These examples demonstrate that surprise-related BOLD signal increases may not be straightforwardly attributed to prediction error signalling, and that further work is required to disentangle co-occurring, confounding phenomena. It remains to be determined why fMRI BOLD signals show different patterns to electrophysiological measures under equivalent conditions.

The third possibility is that cued expectations simply do not influence stimulus-evoked activity of visual stimulus-selective neurons in predictive cueing designs. This would suggest that prior observations of ES and surprise effects are not due to reduced prediction error signalling that occurs when expectations are fulfilled, but are rather due to co-occurring phenomena (such as pupil dilation) or confounding factors (such as adaptation, discussed in Feuerriegel et al., 2021a).

To be clear, we acknowledge that there are consequences of learned expectations that, in certain circumstances, can influence activity in the visual system as well as behaviour. There are documented effects of directed attention guided by expectations (Corbetta et al., 1990; Reynolds & Heeger, 2009) and pre-activation of expected stimulus representations (Kok et al., 2017; Blom et al., 2020; Feuerriegel et al., 2021c). Beyond the visual system, biases in motor action preparation (de Lange et al., 2013; Gold & Stocker, 2017; Kelly et al., 2021) can enable faster responses when expectations to see a particular stimulus are paired with expectations to make a certain motor action. There are also well-documented effects associated with reward predictions errors and information gain in areas such as the insula and striatum (e.g., Loued-Khenissi & Preuschoff, 2020; Schultz, 2016). Rather, our findings suggest that appeals to ES and cortical prediction error minimisation via top-down inhibition are not necessary to adequately model expectation-related phenomena in the visual system. This highlights the utility of alternative models that describe a range of cued expectation effects without recourse to ES (e.g., Spratling, 2017; Heeger, 2017; Press et al., 2020, 2022; Feuerriegel et al., 2021c; Hogendoorn, 2022). Notably, many of these models describe top-down excitatory rather than inhibitory modulations of stimulus-selective visual neurons. This is consistent with evidence that cortical feedback projections within the visual system predominantly originate from, and target, excitatory rather than inhibitory neurons (reviewed in Spratling, 2017).

Mechanisms described in these models are likely to interact with other processes that exploit the spatiotemporal structure of sensory environments. These processes include adaptation (Solomon & Kohn, 2014; Vogels, 2016; Whitmire & Stanley, 2016), surround suppression (Carandini & Heeger, 2012), end-stopping (Rao & Ballard, 1999), as well as retinal circuit mechanisms that rapidly adjust to local luminance levels (Rieke & Rudd, 2009), environmental image structure (Hosoya et al., 2005), periodic stimulation patterns (Schwartz et al., 2007; Schwartz & Berry, 2008) and smooth motion (Berry et al., 1999; Johnson et al., 2023). Systematically investigating these interactions will likely uncover a host of novel predictive phenomena, including feedforward or inherited effects across the visual cortical hierarchy (e.g., Dhruv et al., 2011; Larsson et al., 2016).

### 4.2. Stimulus Predictability Effects

We also tested for stimulus predictability effects, operationalised here as differences in the number of different stimuli that an observer could expect to appear. In Feuerriegel et al. (2018a) we observed predictability effects at electrodes over visual cortex. However, the relatively unpredictable faces in Feuerriegel et al. (2018a) were also more novel than predictable faces. When controlling for this novelty confound in our experiment, we did not replicate these effects.

Notably, we have observed predictability effects on BOLD signals in the Fusiform Face Area (Rostalski et al., 2020) while controlling for a similar novelty confound in Pajani et al. (2017). This appears to be another example of BOLD signal effects that are not replicated when using electrophysiological measures.

### 4.3. Image Repetition Effects

We observed within– and across-trial repetition effects with similar latencies and topographies to those reported in prior work (e.g., Schweinberger & Neumann, 2016; Feuerriegel et al., 2018a, 2019). Our findings complement existing evidence that repetition effects last over multiple intervening stimulus presentations (Bayliss & Rolls, 1987; Kaliukhovich & Vogels, 2014; Henson, 2016; Fritsche et al., 2022) and highlight the need to strictly control for stimulus repetition and recency when testing for consequences of learned expectations.

Our findings are consistent with work demonstrating distinct and separable repetition and expectation effects (Grotheer & Kovács, 2015; Feuerriegel et al., 2018a; Tang et al., 2018) and cortical adaptation in the absence of ES (Kaliukhovich & Vogels, 2011, 2014; Vinken & Vogels, 2018; Solomon et al., 2021). This body of evidence is not congruent with models that describe repetition suppression and adaptation as reflecting biased expectations toward recently seen stimuli (Summerfield et al., 2008; Auksztulewicz & Friston, 2016). These findings instead preferentially support normalisation-based models of adaptation (e.g., Dhruv et al., 2014; Solomon & Kohn, 2014; Vogels, 2016), which can account for a wide range of adaptation-related phenomena without recourse to expectations or prediction error minimisation (e.g., Dhruv et al., 2011; Patterson et al., 2013; Kaliukhovich & Vogels, 2016).

Here, we note that the S1/S2 face repetition effect spanning 379-453ms from S2 face onset resembles the topography of the centro-parietal positivity (CPP) component (O’Connell et al., 2012) and is likely a consequence of faster RTs (and earlier CPP onsets) for repeated as compared to unrepeated stimuli (discussed in Feuerriegel et al., 2022).

### 4.4. Limitations

Our findings should be interpreted with the following caveats in mind. Firstly, the models and empirical evidence discussed here pertain to the visual system and may not be representative of other sensory systems. For example, expectation effects have been reported in auditory predictive cueing designs (e.g., Todorovic & de Lange, 2012). BOLD signals in other brain areas such as the insula have also been found to scale with quantitative measures of uncertainty and surprise (Preuschoff et al., 2008; Mohr et al., 2010; Loued-Khenissi et al., 2020), although these effects may depend on the task-relevance of expected and surprising stimuli (Richter & de Lange, 2019). The EEG signals measured in our experiment may not be sensitive to neuronal activity within the insula and other deep brain structures such as the striatum. In addition, our inferences specifically relate to consequences of expectations that are formed in advance of stimulus appearance. The term expectation has been used to describe other phenomena within predictive coding models (e.g., Rao & Ballard, 1999; see Andersen & Chemero, 2013), which may indeed arise via prediction error minimisation mechanisms (for approximations of predictive coding in the retina see Hosoya et al., 2005; Rieke & Rudd, 2009; Spratling, 2017). Our findings are not directly relevant to those phenomena.

We also have focused on evidence from predictive cueing designs in which there is a clear logic regarding how participants’ expectations are manipulated. Effects attributed to expectation violations have been reported in visual oddball designs, although they are not always consistent across experiments using different stimulus categories (e.g., Kimura et al., 2009; Male et al., 2020; Amado & Kovács, 2016; Feuerriegel et al., 2021b) or tasks (File & Czigler, 2019; Petro et al., 2023). In macaques, ES and surprise effects on electrophysiological measures have been observed in statistical learning designs that involve weeks of training and hundreds of image pairings (e.g., Meyer & Olson, 2011; Ramachandran et al., 2017; Kaposvari et al., 2018). However, such effects have not been consistently replicated across studies with human participants and shorter training periods that more closely match hypothesised timescales of internal model updating in predictive coding accounts (e.g., Manahova et al., 2018; Zhou et al., 2020). The qualitatively different patterns of effects across paradigms highlights the importance of building a coherent body of evidence *within* each paradigm and warrant caution when comparing our findings to those from other designs.

In addition, we manipulated expectations relating to face images, meaning that we cannot rule out ES or surprise effects on neurons selective for other stimulus features (but see Kok et al., 2017; Solomon et al., 2021). Future work could adapt our design to manipulate expectations relating to a broader set of stimulus features.

We also note that our analyses of ERP amplitudes focus on phase-locked EEG responses, rather than non phase-locked effects that may be uncovered using time-frequency analyses (e.g., Zhou et al., 2020; but see Vinken & Vogels, 2018; Solomon et al., 2021). Our experiment was not designed to cleanly isolate time-frequency responses evoked by single stimuli within each trial.

### 4.5. Conclusion

Our findings add to the growing body of evidence showing that expectations in predictive cueing designs do not modulate electrophysiological responses in visual stimulus-selective neurons. This evidence does not support predictive coding-based accounts of how expectations shape neural activity in the visual system. We further show that stimulus repetition and expectation effects are distinct and separable, contrary to predictive coding-based models of repetition suppression and adaptation.

## Acknowledgements

This project was supported by an Australian Research Council Discovery Early Career Researcher Award to D.F. (ARC DE220101508). Funding sources had no role in study design, data collection, analysis or interpretation of results. We thank the research groups led by Floris de Lange, Clare Press and Daniel Yon for their insightful comments on our findings.

