## Supplementary Material for "Stimulus expectations do not modulate visual event-related potentials in probabilistic cueing designs"

Melbourne School of Psychological Sciences. Redmond Barry Building, The University of Melbourne, 3010, Australia

### Numbers of Retained Epochs of EEG Data Per Condition

**Supplementary Table S1.** *Numbers of retained EEG epochs for each condition*

| | S1 Faces by Appearance Probability | | | | S1 Faces by Previous Trial S2 Face | | S2 Faces by S1/S2 Repetition / Alternation | | Cues by Previous Trial Cue | |
|---|---|---|---|---|---|---|---|---|---|---|
| | 90% | 50% | 25% | 10% | Rep | Alt | Rep | Alt | Rep | Alt |
| Mean | 404 | 225 | 113 | 45 | 194 | 583 | 395 | 393 | 109 | 663 |
| Median | 410 | 230 | 115 | 46 | 196 | 589 | 399 | 399 | 111 | 670 |
| SD | 21 | 13 | 6 | 3 | 17 | 31 | 21 | 21 | 11 | 38 |
| Min | 333 | 176 | 88 | 39 | 140 | 486 | 311 | 318 | 80 | 538 |
| Max | 431 | 240 | 120 | 48 | 227 | 629 | 419 | 419 | 132 | 718 |

*Note.* Only trials with correct or erroneous responses to S2 faces were included in analyses. Rep = Repetition. Alt = Alternation. S1 and S2 refer to the first and second face images presented in each trial.

### Results of Mass-Univariate Event-Related Potential (ERP) Analyses Using the Occipital Region of Interest

#### Expectation and Predictability Effects on S1 Face-Evoked ERPs

To provide better coverage of visual cortex beyond our occipito-parietal region of interest (ROI), we additionally performed mass-univariate comparisons across expectancy conditions using ERPs averaged over electrodes Oz, O1, O2, Iz and POz which comprised our occipital ROI. We did not observe statistically-significant differences across any of the compared expectancy conditions after correction for multiple tests. Group-averaged ERPs for each set of conditions, difference waves, standardised Cohen's d effect size estimates (Cohen, 1988) and Bayes factors in favour of the alternative hypothesis are displayed in Supplementary Figure S1. We did not observe differences between ERPs evoked by expected and surprising faces (Supplementary Figure S1A), expected and 50% neutral faces (Supplementary Figure S1B), expected and 25% neutral faces (Supplementary Figure S1C), or 50% neutral and surprising faces (Supplementary Figure S1D). Amplitude differences and standardised effect size point estimates were very small. Bayes factors generally provided moderate evidence in favour of the null (values smaller than 1/3) across the time-course of the S1 face-evoked response.

We also compared ERPs across 50% and 25% neutral conditions to test for predictability effects. Statistically-significant differences were not observed after correction for multiple comparison and Bayes factors were in favour of the null hypothesis across most of the peristimulus time window (Supplementary Figure S1E).

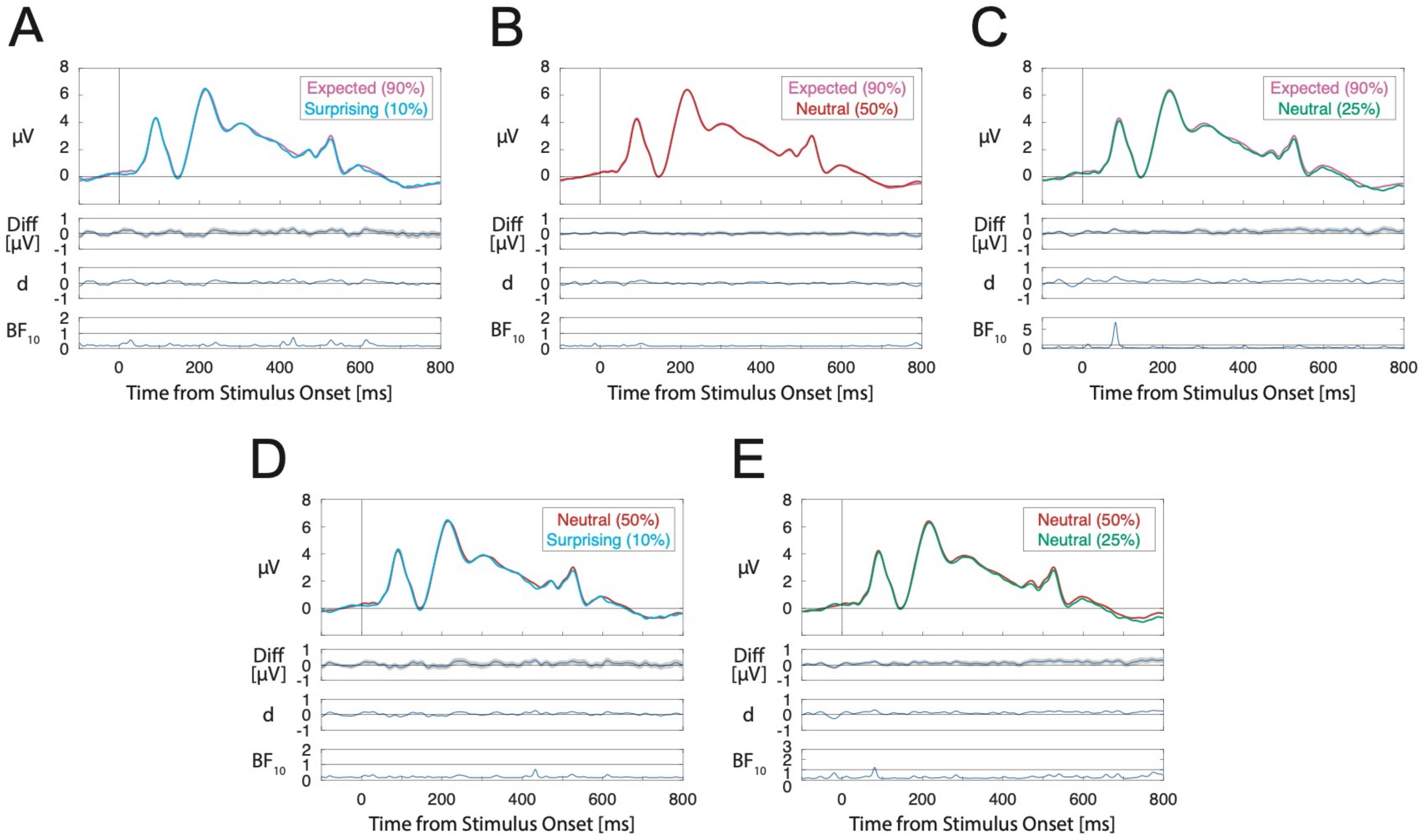
A
B
C
D
E
μV
Expected (90%)
Surprising (10%)
Expected (90%)
Neutral (50%)
Expected (90%)
Neutral (25%)
Neutral (50%)
Surprising (10%)
Neutral (50%)
Neutral (25%)
Diff [μV]
d
$BF_{10}$
0
200
400
600
800
Time from Stimulus Onset [ms]

**Supplementary Figure S1.** Group-averaged ERPs evoked by expected (90% appearance probability), surprising (10% probability), and neutral (50% and 25% probability) S1 faces averaged across channels within the occipital ROI. A) Expected - surprising ERP differences. B) Expected – 50% neutral differences. C) Expected – 25% neutral differences. D) 50% neutral - surprising differences. E) 50% neutral – 25% neutral differences. ERPs for each pair of compared conditions are displayed along with difference waves (with shading denoting standard errors), Cohen's d estimates and Bayes factors in favour of the alternative hypothesis.

**Effects of Within-Trial Repetition on S2 Face-Evoked ERPs**

Multiple, distinct within-trial face repetition effects were observed between 254-800ms from S2 face onset. Group-averaged ERPs for repeated and alternating S2 faces are displayed in Supplementary Figure S2A. This included an effect between 254-332ms (cluster mass = 209.24, critical cluster mass = 50.47, $p = .004$), an effect spanning 357-467ms (cluster mass = 296.10, $p < .001$) and an effect spanning 750-800ms from S2 face onset (cluster mass = 95.17, $p = .033$).

**3.5. Effects of Across-Trial Repetition on Cue- and S1 Face-Evoked ERPs**

Group–averaged ERPs evoked by cues that were the same or different to the cue presented in the preceding trial are shown in Supplementary Figure S2B. No statistically significant across-trial cue repetition effects were observed.

We also compared S1 face-evoked ERPs across conditions whereby the S2 face in the preceding trial was the same or a different face identity. Group-averaged ERPs are displayed in Supplementary Figure S2C. The across-trial repetition effect was observed between 197-232ms (cluster mass = 57.24, critical cluster mass = 36.45, $p = .047$).

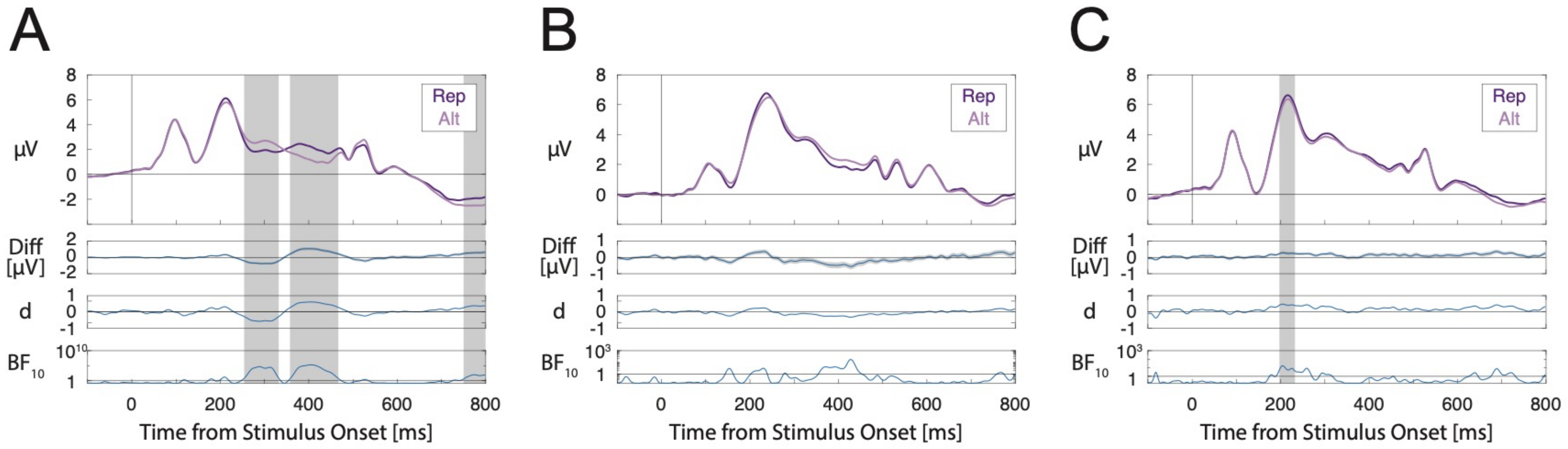


**Supplementary Figure S2.** Within- and across-trial repetition effects for S2 faces, cues, and S1 faces averaged across channels within the occipital ROI. A) S2 face-evoked ERPs depending on whether the S2 face was the same as the S1 face identity (repetition/Rep) or a different face (alternation/Alt). B) Cue-evoked ERPs depending on whether the cue image was the same as the cue in the previous trial (Rep) or a different cue (Alt). C) S1 face-evoked ERPs depending on whether the S2 face in the previous trial was the same (Rep) or a different identity (Alt). ERPs for each pair of compared conditions are displayed along with difference waves (with shading denoting standard errors), Cohen's d estimates and Bayes factors in favour of the alternative hypothesis. Please note that Bayes factors are plotted on logarithmic scales due to the wide ranges of values across the time-course of the stimulus-evoked response. Grey shaded areas denote time windows of statistically significant differences.
